# Nitrogen regulator GlnR directly controls transcription of *prpDBC* operon involved in methylcitrate cycle in *Mycobacterium smegmatis*

**DOI:** 10.1101/353219

**Authors:** Xin-Xin Liu, Wei-Bing Liu, Meng-Jia Shen, Bang-Ce Ye

**Author notes:** These authors contributed equally (Xin-Xin Liu, Wei-Bing Liu).

## Abstract

*Mycobacterium tuberculosis* utilizes the fatty acids of the host as the carbon source. While the metabolism of odd chain fatty acids produces propionyl-CoA. Methylcitrate cycle is essential for Mycobacteria to utilize the propionyl-CoA to persist and grow on these fatty acids. In *M. smegmatis*, methylcitrate synthase, methylcitrate dehydratase, and methylisocitrate lyase involved in methylcitrate cycle were respectively encoded by *prpC*, *prpD*, *and prpB* in operon *prpDBC*. In this study, we found that the nitrogen regulator GlnR directly binds to the promoter region of *prpDBC* operon and inhibits its transcription. The typical binding sequence of GlnR was identified by bioinformatics analysis and electrophoretic mobility shift assay. The GlnR-binding motif was seperated by 164 bp with the binding site of PrpR which was a pathway-specific transcriptional activator of methylcitrate cycle. Moreover, the affinity constant of GlnR was much stronger than that of PrpR to *prpDBC*. The deletion of *glnR* resulted in poor growth in propionate or cholesterol medium comparing with wild-type strain. The Δ*glnR* mutant strain also showed a higher survival in macrophages. These results illustrated that the nitrogen regulator GlnR regulated methylcitrate cycle through directly repressing the transcription of *prpDBC* operon. The finding reveals an unprecedented link between nitrogen metabolism and methylcitrate pathway, and provides a potential application for controlling populations of pathogenic mycobacteria.

**Author Summary:** Nutrients are crucial for the survival and pathogenicity of *Mycobacterium tuberculosis*. The success of this pathogen survival in macrophage due to its ability to assimilate fatty acids and cholesterol from host. The cholesterol and fatty acids are catabolized via β-oxidation to generate propionyl-CoA, which is then mainly metabolized via the methylcitrate cycle. The assimilation of propionyl-CoA needs to be tightly regulated to prevent its accumulation and alleviate toxicity in cell. Here, we identified a new regulator GlnR (the nitrogen transcriptional regulator) that repressed the transcription of *prp* operon involved in methylcitrate cycle in *M. smegmatis*. In this study, we found a typical GlnR binding box in *prp* operon, and the affinity is much stronger than that of PrpR which is known as a pathway-specific transcriptional activator of methylcitrate cycle. In addition, deletion of *glnR* obviously affect the growth of mutant in propionate or cholesterol medium, and show a better viability in macrophage. The findings not only provide the insights into the regulatory mechanism underlying crosstalk of nitrogen metabolism and carbon metabolism, but also reveal a potential application for controlling populations of pathogenic mycobacteria.

## Introduction

Tuberculosis is a chronic bacterial infection which infects one-third of the human population and causes two million deaths annually [1, 2]. Recent researches indicated that fatty acids and cholesterol are more favored carbon sources for *Mycobacterium tuberculosis* when infected in animals and grew within macrophages [1–4]. β-oxidation of even-chain fatty acids can yield acetyl-CoA, while odd-chain fatty acids yield propionyl-CoA as an additional product [5]. Degradation of cholesterol also produces acetyl-CoA and propionyl-CoA [6]. The further metabolism of acetyl-CoA and propionyl-CoA needs glyoxylate and methylcitrate cycle, respectively, in mycobacteria. Methylcitrate cycle converts propionyl-CoA to pyruvate at a molar of 1:1 [7–13]. In *M. tuberculosis*, methylcitrate cycle is not only essential to grow on odd-chain fatty acids but also essential for them to grow in murine bone marrow-derived macrophages. Thus, methylcitrate cycle plays an important role for *M. tuberculosis* to grow, persist, and keep virulence.

Methylcitrate dehydratase (MCD, encoded by *prpD*), methylisocitrate lyase (MCL, encoded by *prpB*), methylcitrate synthase (MCS, encoded by *prpC*) are specific to methylcitrate cycle and essential for mycobacteria to grow on propionate as the sole carbon source [3, 7, 8, 10–12, 14]. *M. tuberculosis* contains only MCD and MCS (encoded by operon *prpDC*) [15–17], and the activity of MCL is provided by isocitrate lyases (ICL1 and ICL2) which are two isoforms involved in glyoxylate pathway. *M. smegmatis* contains *prpDBC* operon encoding three enzymes involved in methylcitrate cycle (*msmeg_6645, msmeg_6646*, and *msmeg_6647*). *PrpD(B)C* operon plays a key role in assimilation of propionyl-CoA for both *M. tuberculosis* and *M. smegmatis* to obtain carbon, energy and prevent the accumulation of toxic metabolite [5, 18]. However, so far the regulation mechanism underlying methylcitrate cycle and *prpD(B)C* operon is little known. PrpR, a transcriptional regulator was reported to regulate methylcitrate pathway by activating *prp* operon in *M. tuberculosis*, and the *prpR* knockout strain exhibited an damaged growth [3]. In *Salmonella enterica Serovar Typhimurium*, PrpR activated *prp* operon in responsive to 2-methylcitrate [19].

Nitrogen plays an important part in bacteria growth and biological macromolecules composition. An OmpR-type transcriptional regulator, GlnR has reported to mediate the expression of more than 680 genes including MEMEG_6645 and MSMEG_6647 involved in methylcitrate cycle [20]. In this study, a typical nitrogen regulator GlnR binding motif (<u>**GGACC**</u>GGCACC<u>**GTAAC**</u>) was observed at the upstream region of *prpDBC* operon in *M. smegmatis*. We found that GlnR directly binds to this region and represses its transcription. The finding reveals an unprecedented link between nitrogen metabolism and propional-CoA assimilation involved in the utilization of fatty acids or cholesterol, and provides a new insight to the sensing and metabolism of nutrients of mycobacteria in host cells.

## Experimental procedures

### Bacterial Strains and Culture Conditions

All bacterial strains and plasmids used in this experiment are listed in Table 1. *Mycobacterium smegmatis* strains were cultured in in LB broth (supplemented with 0.05% Tween 80) at 37 °C, 200 rpm or LB agar plates at 37 °C. For growth on different carbon sources, strains were grown in M9 media containing 12.6 mM Na_2_HPO_4_, 22 mM KH_2_PO_4_, 8 mM NaCl, 19 mM NH_4_Cl, 2 mM MgSO_4_, 0.1 mM CaCl_2_ and 0.1% tyloxapol. Provided carbon sources included 10 mM propionate or 10 mM cholesterol, or other odd-chain fatty acids. All *E. coli* strains used in this experiment were grown in LB broth at 37 °C, 200 rpm.

**Table 1.** Strains and plasmids used in this study

| Bacterial strains and plasmids | Source or reference |
| --- | --- |
| **Bacterial strains** |  |
| *Mycobacterium smegmatis* mc2 155 | [21] |
| *Mycobacterium smegmatis* mc2 155 Δ*glnR* | [21] |
| *Mycobacterium smegmatis* mc2 155 Δ*glnR::glnR* | In this study |
| *E. coli* DH5α | Novagen |
| *E. coli* BL21(DE3) | Novagen |
| **plasmid** |  |
| pET-28a | Novagen |
| pET-28a-*glnR* | In this study |
| pET-28a-*prpR* | In this study |

### Cloning, overexpression, and purification of GlnR and PrpR

The genes *glnR* and *prpR* were amplified by PCR from the genome of *M. smegmatis* mc2 155. Seamless cloning and assembly kit was used. After purification, the PCR product was introduced into pET-28a to generate recombinant vector, pET-28a-*glnR* or pET-28a-*prpR*. The clones were confirmed by PCR and sequencing. The proteins were expressed by *E. coli BL21*(DE3). A single clone was cultured in 5 ml LB (1‰ kanamycin) overnight, and then transferred to 50 ml LB (1‰ kanamycin). 0.5 mM IPTG was added when the OD_600_ of the cells was about 0.7. Then the cells were grown at 20°C overnight.

Cells were collected by centrifugation at 8000 g for 10 min then re-suspended in 25 ml PBS buffer. The cells were disrupted using sonication, and the cell debris was removed by centrifugation. The supernatant was purified using Ni-NTA agarose column (Merck) that was pre-equilibrated in 10 mM imidazole in 50 mM NaH_2_PO_4_ and 300 mM NaCl, pH 8.0. The desired protein was eluted with 20- 250 mM imidazole in 50 mM NaH_2_PO_4_ and 300 mM NaCl (pH 8.0). The fractions were analysized by SDS-PAGE electrophoresis. The protein concentration was determined by the BCA method.

### Electrophoretic mobility shift assay (EMSA)

The upstream region (from −300 to 50) of *prpDBC* containing GlnR-binding and PrpR-binding sites were amplified by PCR with biotin-labeled primer (5’ biotin- AGCCAGTGGCGATAAG 3’). The PCR product was analyzed by agarose gel electrophoresis, and purified using a PCR Purification kit (Shanghai Generay Biotech). The concentrations of biotin-labeled DNA probes were determined with a microplate reader (Biotek, USA). EMSAs using N-terminally His-tagged GlnR or PrpR were carried out following the Chemiluminescent EMSA Kit (Beyotime Biotechnology, China) manual. The binding mixture (total volume 10 μl) containing 1 μl DNA probes, varying amounts of purified GlnR or PrpR and 1 μl Gel-Shift binding buffer was incubated at room temperature for 20 min. After binding, the mixture was separated on a non-denaturating PAGE gel in ice-bathed 0.5× Tris-borate-EDTA at 100 V and bands are detected by BeyoECL Plus.

### Quantitative real-time PCR

Cells at exponential stage in nitrogen-limited medium were collected by centrifugation. Total RNA was prepared using Rneasy Mini Kit (Qiagen, Valencia, CA). The RNA quality was analyzed by 1% agarose gel electrophoresis and the concentration was determined by microplate reader (BioTek, USA). The RNA was reverse transcribed to cDNA using a PrimeScript™ RT Reagent Kit with gDNA Eraser (Takara, Shiga, Japan), and DNase digestion was performed to remove genomic DNA before reverse transcription for 5 min at 42°C. PCR reactions were performed with primers listed in Table 2.

Real-time PCR reactions was performed using 2 × RealStar Green Fast Mixture (GeneStar, Beijing, China) in 20 μL final volume containing 50 ng cDNA on a CFX96 Real-Time System (Bio-Rad, USA). The PCR reaction conditions were as follows: 95°C for 5 min, 40 cycles of 95°C for 5 s and 60°C for 30 s, then extended at 72°C for 10 min. For all the RT-PCR assays, 16s RNA was used as an internal control. The transcriptional fold changes of target genes were calculated using 2^^-ΔΔCt^ method.

**Table2.**
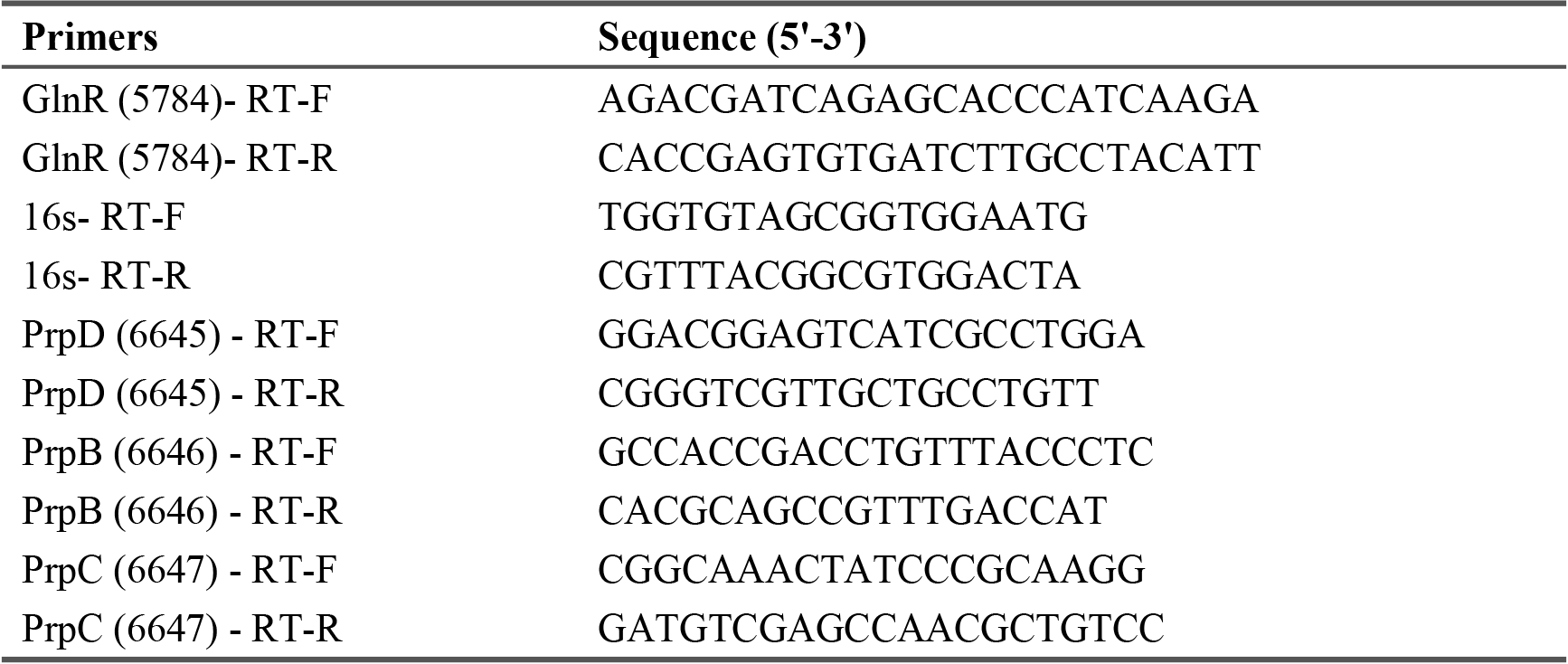
The primers used for real-time PCR

### Kinetic binding analysis using Octet system

The binding affinities of GlnR and PrpR proteins to the upstream region of *prpDBC* operon were determined by Bio-Layer interferometry using an Octet System (Octet Q^Ke^, Fortebio, USA). Streptavidin biosensors were loaded with biotinylated DNA fragment (upstream region of *prpDBC)* by incubation for 5 min in 7 ug/ml DNA solution and then washed in loading buffer for 5 min. After that, the biosensors were moved to protein solutions to allow association for 10 min and then transferred into PBS buffer to detect dissociation. All samples were performed in a 96-well plate at 37°C and 1000 rpm in a volume of 100 μl. The proteins were diluted in the PBS buffer containing 10% glycerol, 10 μg/ml BSA, and 0.02% tween-20. The kinetic parameters-k_on_, k_off_, and K_D_ were calculated by 1:1 binding model using the Octed Data Analysis version 7.0.

### Resazurin assays

To monitor the cell metabolism activity of three *M. smegmatis* strains (wild-type, *glnR*-deleted mutant Δ*glnR*, and complementary strain Δ*glnR::glnR*) on different carbon sources, the resazurin assay was performed. Strains were activated by culturing in LB containing 0.05% tween-80, and then transferred to M9 media with different carbon resources at the same initial OD_600_. After 36 h of growth, 100 μl of the culture was added into 96-well plate. Then resazurin solution (12.5 mg/ml final concentration, Sigma) was added into the plate. The change in color was observed every ten minutes. Media without cells and cells without carbon source were used as negative controls [22].

### Survival of M. smegmatis in macrophages

The human mononuclear macrophage THP-1 cell was from the Cell Bank of Typical Culture Preservation Committee of Chinese Academy of Science (Shanghai, China), and was cultured in RPMI-1640 medium with 10% FBS at 37 °C in the presence of 5% CO_2_. They were divided into 12-well plates (2.0 × 10^5^ cells per well) and differentiated using PMA when the cells were at the best state [23]. After 12 h, the cells were washed and then cultured in fresh RPMI-1640 medium for 12 h. The *M. smegmatis* were added into the plates (ten times more than the THP-1 cells). After incubated for 2 h, the cells were washed three times using fresh medium to remove the uninfected bacteria. The infected cells were cultured in fresh medium with gentamicin. At the different infected time, the cells were washed three times with fresh medium, and then added LB medium containing 0.05% SDS to lyse the cells for 10 min. The lysates were collected and diluted at different gradient to inoculate plate. The *M. smegmatis* Colony-Forming Units were counted after 3 days culturing [24].

## Results

### Nitrogen response regulator GlnR binds the promoter region of prpDBC operon

In previous work, we found that nitrogen regulator GlnR in actinobacteria directly regulated carbon metabolisms, including the uptake and utilization of non-phosphotransferase-system carbon sources [25], degradation of starch [26], synthesis of citrate [27], and synthesis of acetyl-CoA/propionyl-CoA [24, 28]. In this study, a typical GlnR-binding motif (<u>**GGACC**</u>-GGCACC-<u>**GTAAC**</u>) was identified in the upstream region of *prpDBC* operon involved in methylcitrate cycle in *M. smegmatis* (Fig 1A). To examine whether GlnR can bind this predicted motif or not, we use diluted different gradient of GlnR protein with DNA probe containing the putative GlnR-binding motif in EMSA assay. 200-fold excess of unlabeled probe (S) and sperm DNA (N) were used as controls. As shown in Fig 1B, obvious shift bands were observed when the purified GlnR were added, and the shift bands increase with GlnR concentrations. The results demonstrated that GlnR was able to directly bind the upstream region of *prpDBC* operon specifically *in vitro*, and suggesting that *prpDBC* may be subjected to transcriptional regulated by the nitrogen regulator GlnR.

**Fig 1.**
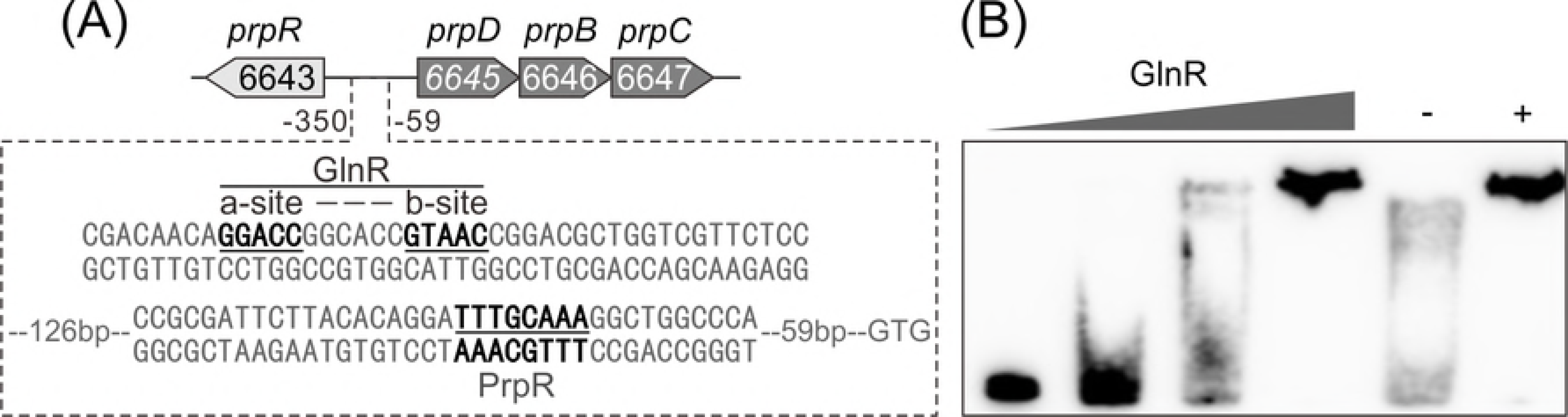
Nitrogen response regulator GlnR binds the promoter region of *prpDBC* in *M. smegmatis*. (**A**) Cis-element analysis of *prpDBC* promoter region. (**B**) EMSA of His-GlnR with regulatory region of *prpDBC*. 10 mM DNA probe was incubated with a gradient concentration of GlnR (0, 0.3, 1.2, 4.8 μM). 200-fold excess of unlabeled probe (S) and non-specific competitor DNA (sperm DNA) (N) were used as controls.

### GlnR repressed the transcription of prpDBC operon in M. smegmatis

To investigate the regulatory effect of GlnR on *prpDBC in vivo*, we constructed *glnR*-deleted strain (Δ*glnR*) and the complementary strain (Δ*glnR::glnR*) of *M. smegmatis*. Three strains (wild-type, Δ*glnR*, and *ΔglnR::glnR)* were cultured in nitrogen-limited M9 medium. The transcriptional levels of *prpD, prpB*, and *prpC* in three strains were compared. As shown in Fig 2, the transcriptional level of *prpDBC* increased obviously in *glnR*-deleted strain compared to the wild type strain. The transcripts were increased 8-fold for *prpD*, 4-fold for *prpB*, and 6-fold for *prpC* in Δ*glnR* strain. The complementation of *glnR* gene into Δ*glnR* strain resulted in recovery of *prpDBC* transcripts (Fig 2). The data showed that GlnR repressed the transcription of *prpD, prpB*, and *prpC in vivo* in *M. smegmatis*.

**Fig 2.**
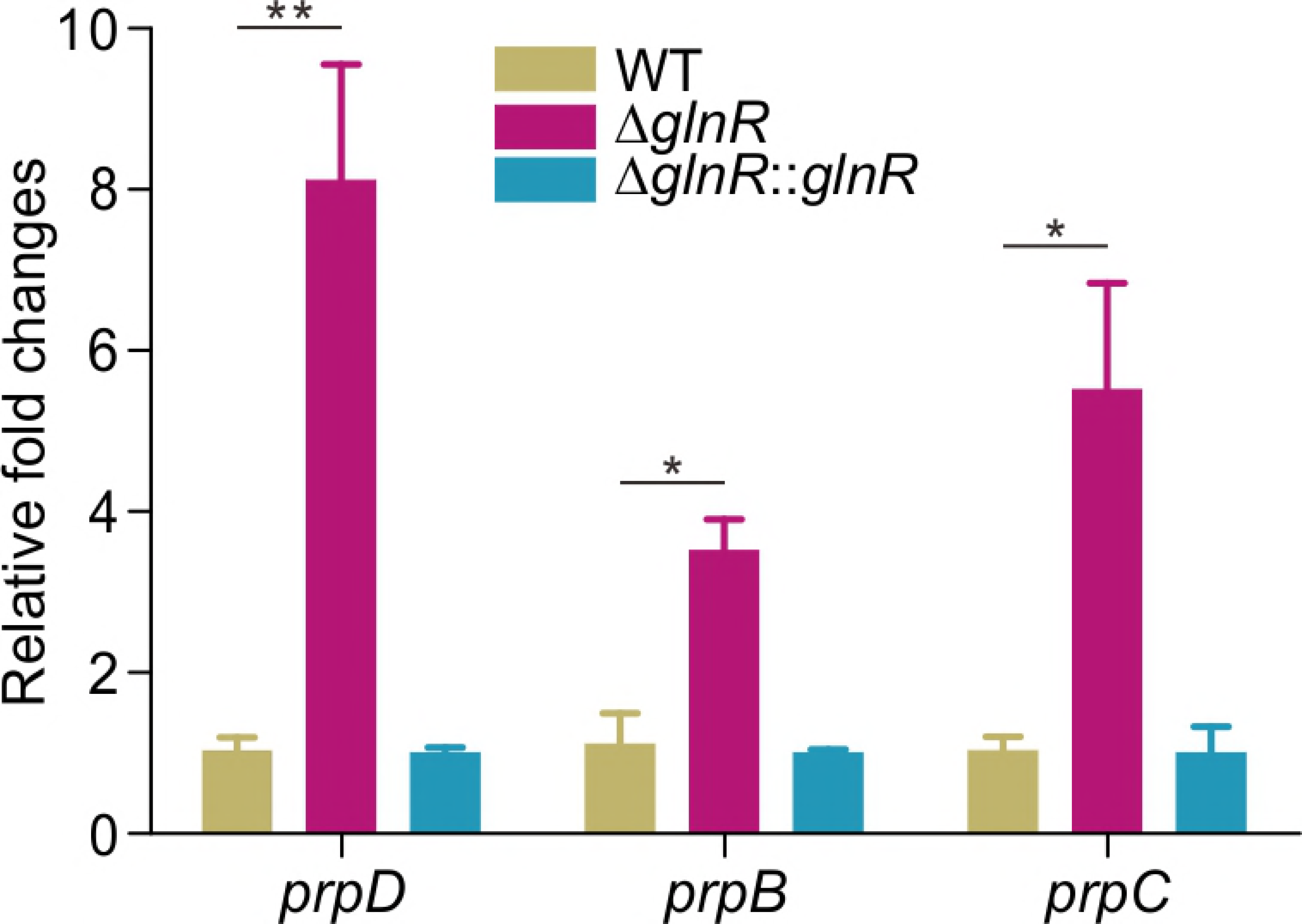
GlnR repressed the transcription of prpDBC in *M. smegmatis*. The transcriptional analysis of prpDBC in *M. smegmatis* wild type strain (WT), glnR-deleted strain (ΔglnR) and complementary strain (Δ*glnR::glnR*) grown in nitrogen starvation M9 medium. Fold changes represents the transcription levels of *prpDBC* compared to that in wild-type strain. Error bars were the standard deviation of three independent experiments. *P<0.01, **P<0.001

### The transcription of prpDBC is responsive to nitrogen availability

GlnR is a global nitrogen regulator that regulates the transcription of the genes involved in nitrogen metabolism is responsive to nitrogen availability [24, 29]. The transcriptional response of *prpDBC* operon was investigated under nitrogen-limited (N^L^) and nitrogen-rich (N^EX^) conditions. As shown in Fig 3, the limitation of nitrogen resulted in a 5-fold increase for *glnR*, but 73% decrease for *prpD*, 65% decrease for *prpB*, and 61% decrease for *prpC*. GlnR-mediated repression of *prpDBC* operon was observed under nitrogen-limited condition. This result further demonstrated that GlnR directly controls transcription of *prpDBC* operon involved in methylcitrate cycle in response to nitrogen availability in *M. smegmatis*.

**Fig 3.**
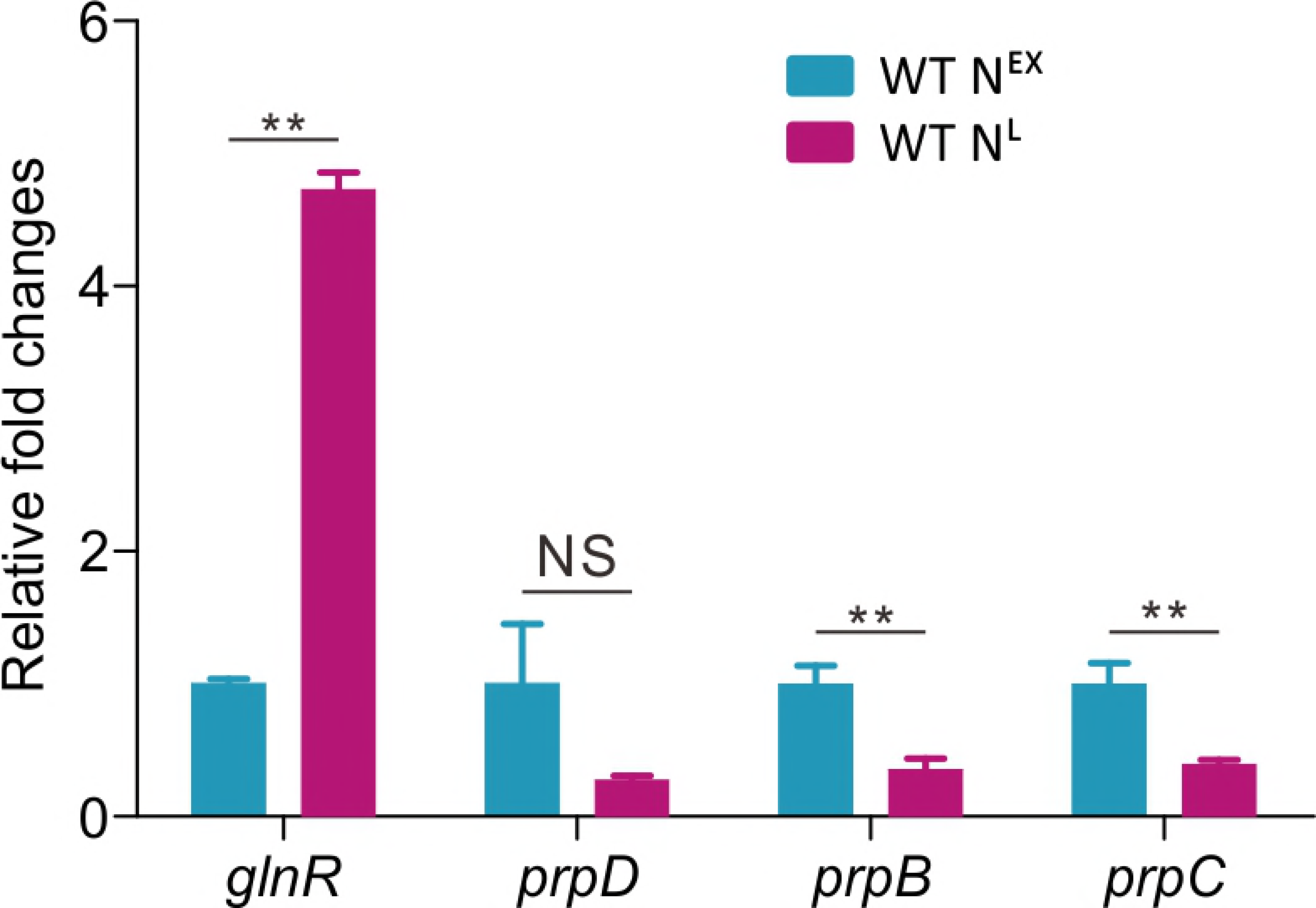
The transcription of *prpDBC* is responsive to nitrogen availability. Wild type strain (WT) was grown in M9 medium with excess (N^EX^) or limited (N^L^) nitrogen. Fold changes represents the transcription levels of *prpDBC* in limited nitrogen compared to that in excess nitrogen. Error bars were the standard deviation of three independent experiments. *P<0.01, **P<0.001.

### GlnR shows higher affinity to prpDBC promoter than PrpR

A typical GlnR binding motif (<u>GGACC</u>-GGCACC-<u>GTAAC</u>) was found in the upstream region of *prpDBC* operon, and confirmed by the EMSA assays. To verify whether the motif sequence is key for GlnR-binding, two biotin-labeled synthetic probes (128 bp) containing the predicted binding motif (P1) and mutant motif (P2) were used for EMSA assays. As shown in Fig 4A and 4B, no shift band was observed for probe P2, indicating that the predicted GlnR-binding sequence in the promoter region of *prpDBC* was directly bound by GlnR. Transcriptional regulator PrpR was reported to directly regulate *prpD(B)C* operon in *M. tuberculosis*, *M. smegmatis*, and *S. enterica*. An 8-bp PrpR-binding motif sequence (TTTGCAAA) was identified in *M. tuberculosis* H37Rv. In this work, a PrpR binding motif (TTTGCAAA) in the upstream region of *prpDBC* was observed, and separated by 164 bp with GlnR binding sequence (Fig 1A). To determine the binding affinities of these two regulators to the upstream region of *prpDBC*, Octet assays were performed using proteins with different concentrations gradients. The KD values of GlnR and PrpR for *prpDBC* are 38.5 and 404 nM, respectively (Fig 4C and 4D). These results revealed that GlnR has a 10-fold higher affinity for the promoter region of *prpDBC* than PrpR.

**Fig 4.**
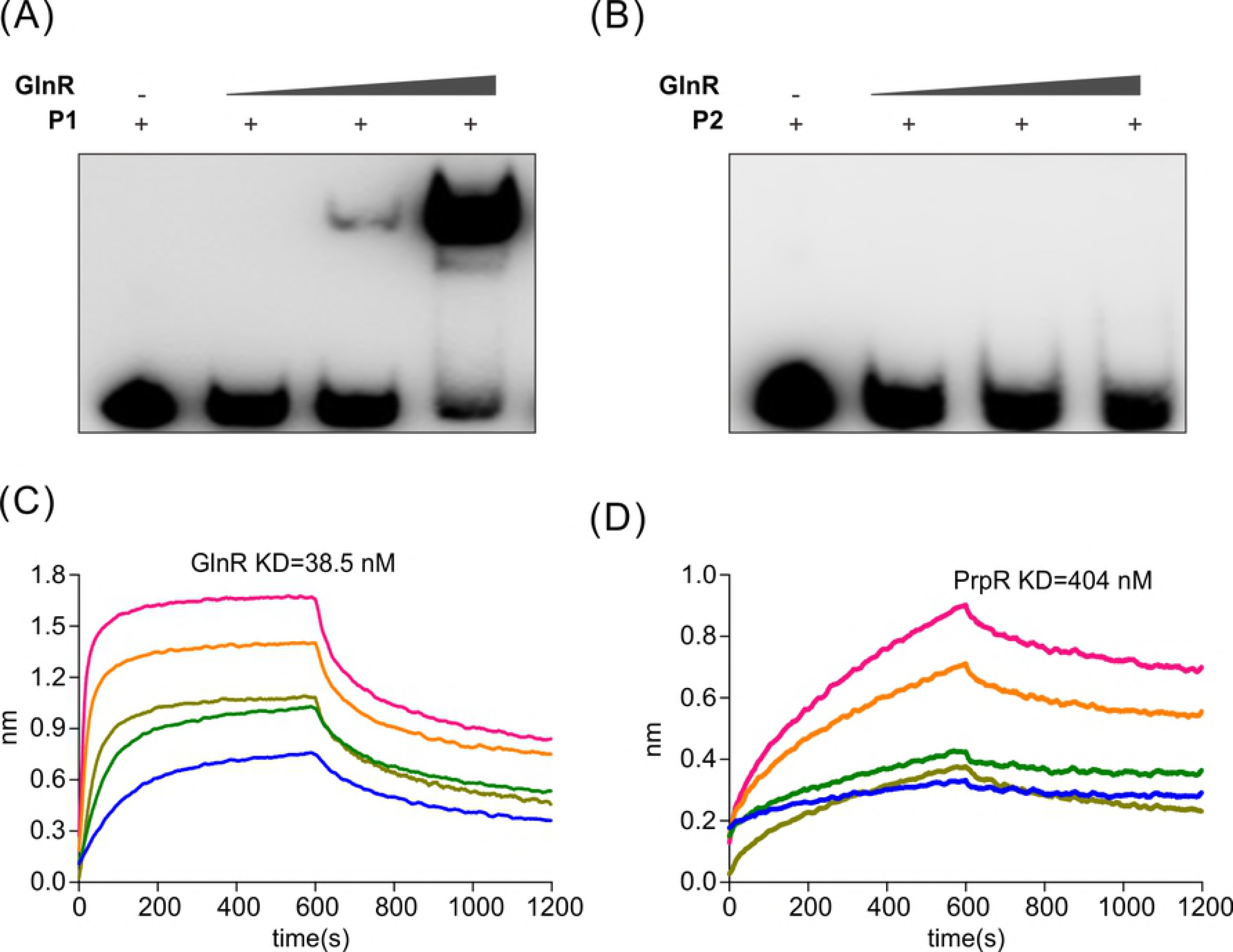
Binding of GlnR or PrpR with the upstream region of *prpDBC*. (A). EMSA of His-GlnR with biotin labeled EMSA fragment (P1). <u>GGACC</u>GGCACC<u>GTAAC;</u> (B). EMSA of His-GlnR with biotin labeled EMSA fragment (P2). <u>**A**GAC**T**</u>GGCACC<u>**A**T**GGT**</u>; (C) and (D). Binding of *prpDBC* biotin labeled probe with increasing concentration of His-GlnR (2.4, 1.2, 0.6, 0.3, 0.015 μM) (C) and His-PrpR (6.4, 3.2, 1.6, 0.8, 0.4, 0 μM) (D).

### GlnR affects the growth of M. smegmatis on propanoate or cholesterol

GlnR repressed the transcription of *prpDBC* operon, which was involved in methylcitrate cycle, and played an important role in metabolisms of fatty acid and cholesterol for mycobacteria in host cells. It is reasonable to expect that GlnR will exert an effect on the growth of *M. smegmatis* on fatty acid or cholesterol as carbon source. To investigate the regulatory effect of GlnR on utilization of fatty acid or cholesterol, three *M. smegmatis* strains (wt, Δ*glnR*, and Δ*glnR::glnR*) were cultivated respectively on 10 mM propanoate or cholesterol. As shown in Fig 5A and 5B, Δ*glnR* strains grew much better than wild-type strain both on 10 mM or cholesterol under nitrogen-limited condition. The deletion of *glnR* alleviated GlnR-mediated repression of *prpDBC* operon, and increases activity of methylcitrate pathway (assimilation of propanoate and propionyl-CoA).

**Fig 5.**
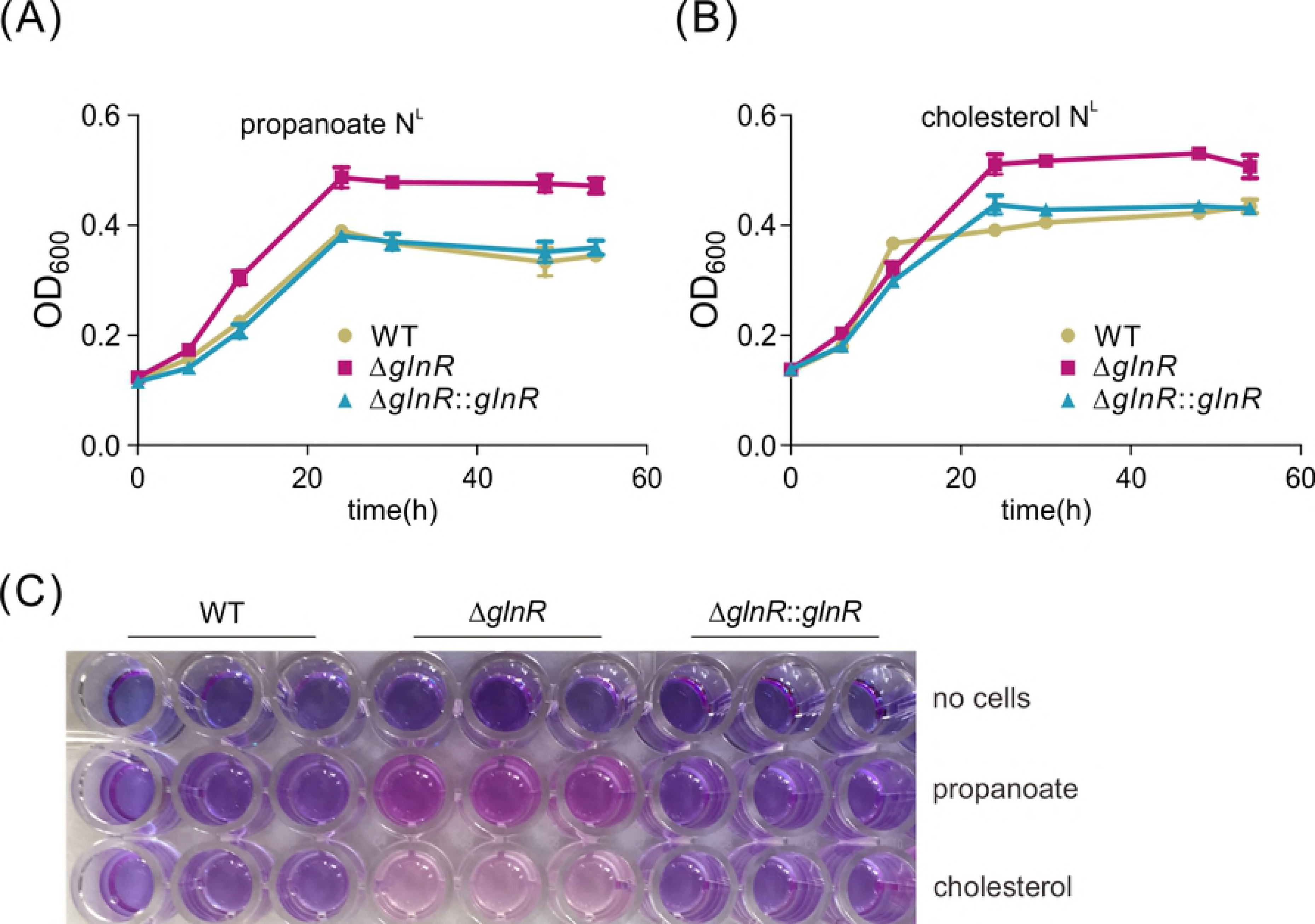
Growth curves of WT, Δ*glnR*, Δ*glnR::glnR* strains in minimal medium with propionate or cholesterol. (A-B). Growth curves of *M. smegmatis* WT, Δ*glnR*, Δ*glnR::glnR* strains on 10 mM propionate (left) or cholesterol (right). (C). Resazurin assay of *M. smegmatis* WT, Δ*glnR*, Δ*glnR::glnR* strains cultured in propionate or cholesterol. Pink represents the cells metabolically active, while blue represents low resazurin reduction and low metabolic activity.

The resazurin reduction assay was also employed to examine the growth of *M. smegmatis* strains on propionate or cholesterol [30]. Blue compound Resazurin, which can be reduced to pink fluorescent product by the metabolically active cells, usually was used to report the cell metabolism activity and cell number. Three *M. smegmatis* strains (wt, Δ*glnR*, and Δ*glnR::glnR*) were grown in nitrogen starvation M9 medium with propionate or cholesterol as carbon source. After 36 h culture, 100 μL of culture were transferred to 96-well plate and resazurin was added. As shown in Fig 5C, Δ*glnR* strain showed a more obvious pink (reduced resazurin) than wild-type and complementary strain. No reduction of resazurin was observed in control wells (no cells). The observations were consistent with the previous conclusion that GlnR repressed the transcription of *prpDBC* and inhibited methylcitrate pathway. The upregulation of *prpDBC* transcription in Δ*glnR* strain resulted in improvement of propional-CoA assimilation for accelerating utilization of propionate or cholesterol.

### GlnR affects the survival of M. smegmatis in macrophages

Lastly, we investigated the effect of GlnR on the survival of *M. smegmatis* in macrophages after infection. Equal amounts of *M. smegmatis* WT, Δ*glnR*, and Δ*glnR::glnR* strains were used to infect the macrophage, and their viability was determined at different time point. As shown in Fig 6, the Δ*glnR* strain revealed the similar survival in macrophages with wild-type strain in the initial 2 h after infection. After 12 h or 24 h, Δ*glnR* strain had a twice survival than wild-type and Δ*glnR::glnR* strains. These results indicated that GlnR exerted a negative effect on the survival of M. *smegmatis* in macrophages. Taken together, GlnR repressed the methylcitrate pathway and propional-CoA assimilation in response to nutrient signals in macrophages, which affected the viability and infection of M. *smegmatis*.

**Fig 6.**
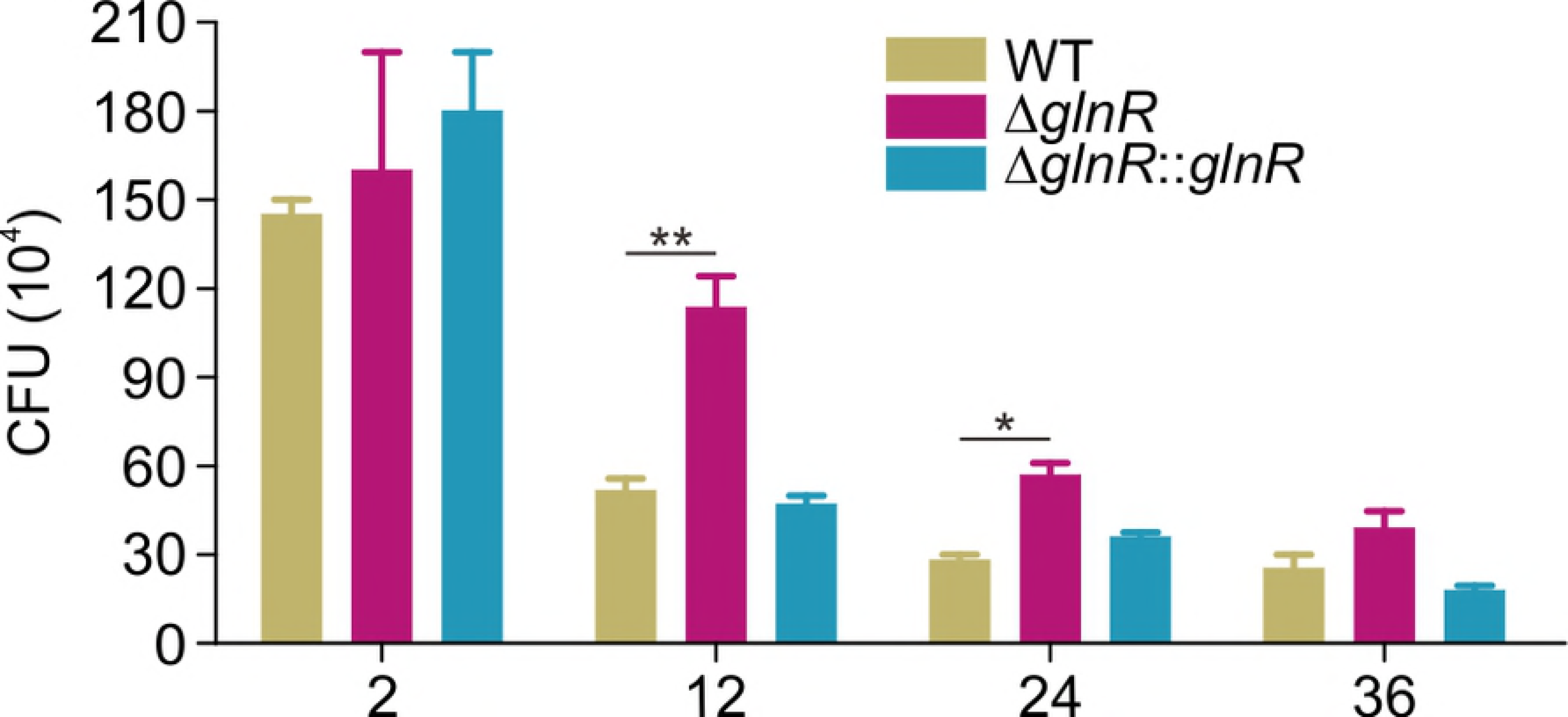
The survival of *M. smegmatis* WT, Δ*glnR*, Δ*glnR::glnR* strains in PMA-differentiated THP-1 macrophages after infected 0-36 h. Data are representative of three independent biological replicates. *P<0.01, **P<0.001

## Discussion

In present study, we identified a new regulator GlnR (the nitrogen transcriptional regulator) that repressed the transcription of *prp* operon involved in methylcitrate cycle in *M. smegmatis*. The finding reveals an unprecedented link between nitrogen metabolism and propional-CoA assimilation involved in the utilization of fatty acids or cholesterol, and provides a new insight to the sensing and metabolism of nutrients of mycobacteria in host cells (Fig 7).

**Fig 7.**
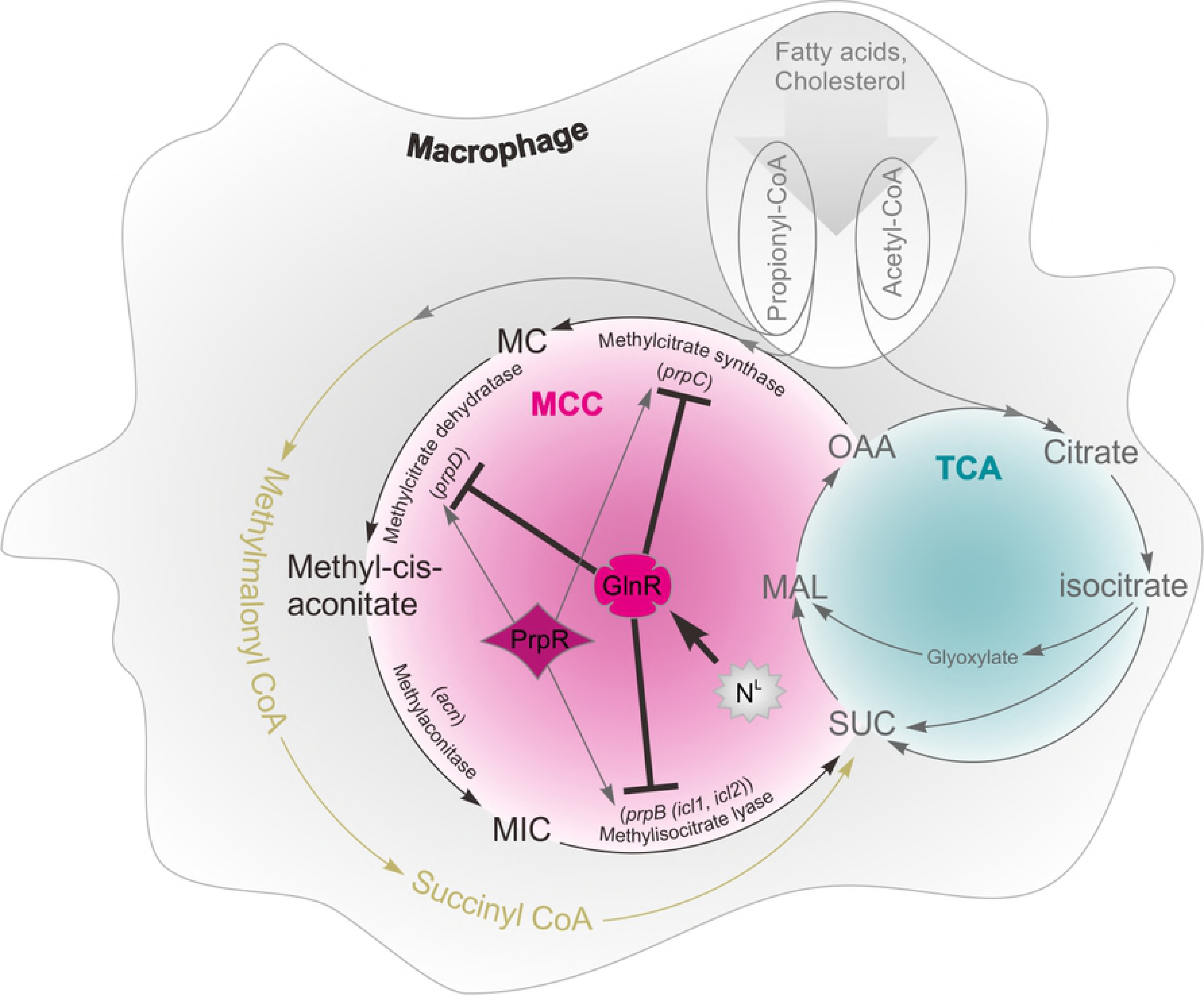
The regulatory mechanism of GlnR on *prpDBC* involved in methylcitrate cycle. MCC, Methylcitrate cycle; MC, Methylcitrate; MIC, Metylisocitrate; SUC, Succinate; MAL, Malate; OAA, Oxaloacetate

Fatty acids and cholesterol metabolism is essential for mycobacteria to grow in macrophages and infect human or animals. However, the degradation of fatty acids and cholesterol results in accumulation of propionyl-CoA [31–33]. Propionyl-CoA is an inhibitor of several key metabolic enzymes such as pyruvate dehydrogenase, succinyl-CoA synthetase and ATP citrate lyase, is considered as a toxic metabolite. Mycobacteria are highly sensitive to increases in the propionyl-CoA pool and it has evolved several different mechanisms for the detoxification of propionyl-CoA or assimilation or propionyl-CoA. So far, two pathways were found for propionyl-CoA assimilation in mycobacteria: the methylcitrate cycle (MCC) that converts propionyl-CoA to pyruvate/succinyl-CoA, and the propionyl-CoA carboxylase (PCC) pathway, which is responsible for the metabolism of propionyl-CoA to methylmalonyl-CoA [33, 34]. These carbon intermediates can be used as precursors participating in the synthesis of pathogenic cell wall lipids. Metylcitrate cycle is the major pathway for propional-CoA utilization for both bacteria and fungi. The assimilation of propionyl-CoA needs to be tightly regulated to prevent its accumulation and alleviate toxicity in cell. The propionyl-CoA assimilation have been investigated in *M. tuberculosis*. It was found that the pathway-specific regulator PrpR regulated methylcitrate cycle by activating *prp* operon in mycobacteria [3, 34]. Previous works have demonstrated that the transcripts of *prp* operon were increased obviously when infected the mouse lung [18, 35], and the Δ*prp* strain wasn’t able to grow and reproduce in murine macrophages [3].

The nitrogen regulator GlnR is activated under condition of nitrogen starvation, and inhibits the assimilation of propional-CoA through directly repressing the transcription of *prpDBC* involved in metylcitrate cycle. The repression of the propional-CoA assimilation results in the poor growth of mycobacteria in propionate or cholesterol. By integrating environmental nitrogen signals to modulate the propional-CoA assimilation involved in the utilization of fatty acids or cholesterol, GlnR mediates the interplay between nitrogen and carbon metabolism of mycobacteria during an infection process. *M. tuberculosis* uses the fatty acids and cholesterol from the host as a carbon source, generating propional-CoA. However, there are seldom reports about the nitrogen metabolism when *M. tuberculosis* grows in host cell. Various nitrogen sources can be used by *M. tuberculosis*, but organic sources (such as amino acids) are more efficient [36], and host- acquired Asp and Asn have been experimentally confirmed as a nitrogen source. This study explains GlnR sensing the nitrogen signal and then regulating propional-CoA assimilation, which may affect virulence lipids synthesis [37].

The findings not only provide the insights into the regulatory mechanism underlying crosstalk of nitrogen metabolism and carbon metabolism, but also reveal a potential application for controlling populations of pathogenic mycobacteria.

